# Contrast Solution Properties and Scan Parameters Influence the Apparent Diffusivity of Computed Tomography Contrast Agents in Articular Cartilage

**DOI:** 10.1101/2021.10.01.462834

**Authors:** Mary E. Hall, Adam S. Wang, Garry E. Gold, Marc E. Levenston

## Abstract

The inability to detect early degenerative changes to the articular cartilage surface that commonly precede bulk osteoarthritic degradation is an obstacle to early disease detection for research or clinical diagnosis. Leveraging a known artifact that blurs tissue boundaries in clinical arthrograms, contrast agent diffusivity can be derived from computed tomography arthrography (CTa) scans. We combined experimental and computational approaches to study protocol variations that may alter the CTa-derived apparent diffusivity. In experimental studies on bovine cartilage explants, we examined how contrast agent dilution and transport direction (absorption vs. desorption) influence the apparent diffusivity of untreated and enzymatically digested cartilage. Using multiphysics simulations, we examined mechanisms underlying experimental observations and the effects of image resolution, scan interval and early scan termination. The apparent diffusivity during absorption decreased with increasing contrast agent concentration by an amount similar to the increase induced by tissue digestion. Models indicated that osmotically induced fluid efflux strongly contributed to the concentration effect. Simulated changes to spatial resolution, scan spacing and total scan time all influenced the apparent diffusivity, indicating the importance of consistent protocols. With careful control of imaging protocols and interpretations guided by transport models, CTa-derived diffusivity offers promise as a biomarker for early degenerative changes.

## INTRODUCTION

Osteoarthritis (OA) is a whole joint disease physically characterized by progressive degradation of the soft tissues and by bony changes including osteophyte formation and subchondral sclerosis and clinically manifest as pain, stiffness and reduced mobility. Symptomatic OA affects approximately 12.1% of the adult population in the United States and costs the health care system an estimated $89.1 billion per year in direct costs^1^, with cases rising due to aging of the population and the increased prevalence of obesity. OA treatments currently aim at managing the condition through pain reduction, improving quality of life, and delaying severe symptoms, as there is no cure^2^ and advanced degeneration is often addressed via total joint replacement. The current standard for radiological diagnosis of knee OA is visible joint space narrowing and the presence of osteophytes on a standing radiograph^3^. By the time such joint space narrowing is present, irreparable damage to the avascular cartilage tissue has occurred^4^. Detection of early degenerative changes linked to OA such as proteoglycan loss^5–8^ or changes in collagen structure of the cartilage^5, 7, 8^ using quantitative imaging techniques could allow for earlier intervention and improved patient outcomes for this disease, as well as studies of pharmaceutical or therapeutic interventions earlier in disease progression.

Although magnetic resonance imaging (MRI) is the modality of choice for more detailed characterization of cartilage lesions in high-income nations, techniques based on computed tomography (CT) have been explored because of high spatial resolution, rapid scan time and relatively low cost. Computed tomography arthrography (CTa) enables visualization of articular cartilage morphology in vivo after intra-articular injection of an iodinated contrast agent to delineate the boundary between soft tissues and the synovial space. While less commonly applied to the knee, CTa has advantages for clinical diagnosis or research studies of other joints including the shoulder^9, 10^, wrist^11, 12^, hip^13^, elbow^14^ and ankle^15^. Over time, penetration of the contrast agent into the soft tissues blurs this boundary and impairs morphological assessment^16, 17^, but various approaches have been explored to leverage contrast agent penetration to quantitatively evaluate cartilage integrity. The distribution of an ionic contrast agent within cartilage is related to the distribution of negatively-charged proteoglycans and the charge of the iodine-bearing ion^18–22^, although this relationship is strongest after equilibration times far longer than is clinically feasible and clinical adoption is limited by the predominant use of non-ionic contrast agents in the United States. Dual^23–26^ and triple^27^ contrast techniques, where cartilage is exposed to multiple contrast agents in order to calculate partition of multiple contrast agents and/or alter the image contrast to allow for easier segmentation, have been used in conjunction with dual energy acquisitions to improve evaluation of proteoglycan content, water content, and morphology at earlier time points. Multiple studies have shown that contrast agent penetration in CT arthrograms is higher in osteoarthritic^28, 29^ or damaged^30, 31^ cartilage than in intact cartilage, motivating studies specifically focused on the kinetics of contrast agent penetration to characterize the physical status of the tissue. Unlike the equilibrium distribution or partition, the diffusivity of solutes in cartilage is predominantly related to the solute size (molar mass or hydrodynamic radius) rather than the solute charge^32^, suggesting that either ionic or non-ionic contrast agents have utility for studying contrast agent kinetics.

Under the assumption that the dominant mode of transport is diffusion, previous studies have calculated the apparent diffusivity of contrast agents in articular cartilage by performing multiple CT scans of cartilage explants over a period of time and fitting models of varying complexity to the average attenuation of the samples^30, 33–37^. Apparent diffusivity could be a promising quantitative biomarker for degradation because it is not directly dependent on the absolute amount of contrast that has penetrated the cartilage at a given scan, but rather is derived from relative differences in contrast distributions between scans. CT-based apparent diffusivity, particularly at early time points, also has the advantage of being derived from phenomena dominated by the properties of the cartilage surface where proteoglycan depletion, mild fibrillation and other early degenerative changes are often present before bulk changes occur^38^. However, most previous analysis approaches relied on idealized assumptions that may deviate substantially from phenomena in actual in vivo scans, and the extent to which these assumptions limit the ability to robustly detect physical differences has not been critically examined.

This could account for the effects on local contrast values of heterogeneous contrast agent distribution in the joint in space and across scan time points that results from heterogeneous initial distribution, contrast pooling over time, osmotically-induced joint effusion and physiologic clearance of the contrast agent from the joint.

When contrast is injected intra-articularly for an arthrogram, it spreads nonuniformly throughout the joint and, over time, a combination of a heterogeneous initial distribution, contrast pooling over time, osmotically-induced joint effusion, and physiologic clearance all influence the concentration at the tissue-fluid interface, and initial soft tissue absorption of the contrast agent may be followed by desorption as the external concentration drops. In addition, use of different types of CT scanners and different scan protocols across research groups can make comparison of results difficult. If quantitative measurements are being made, it is important to understand how all of these factors affect apparent diffusivity calculation. Previous work^35, 39^ has focused on studying effects of concentration in explants over the course of long time scales up to 48 hours, which would not be possible in vivo due to clearance of the contrast agent from the patient’s joint. In this study, we employed a combination of experimental studies on isolated articular cartilage explants and computational simulations to examine effects of a range of physical conditions and imaging parameters on the apparent diffusivity derived from CTa on a short, clinically-relevant timescale. Specifically, experimental studies were used to evaluate interacting effects of contrast agent dilution, imaging during contrast agent absorption versus desorption and experimental tissue degeneration on the derived diffusivity, while simulations were used to explore mechanisms responsible for experimental observations and to investigate the individual effects of spatial resolution, scan frequency and total scan time on the value of the derived diffusivity.

## METHODS

### Experimental Methods

Figure 1 provides an overview of the following experimental procedures.

**Figure 1.**
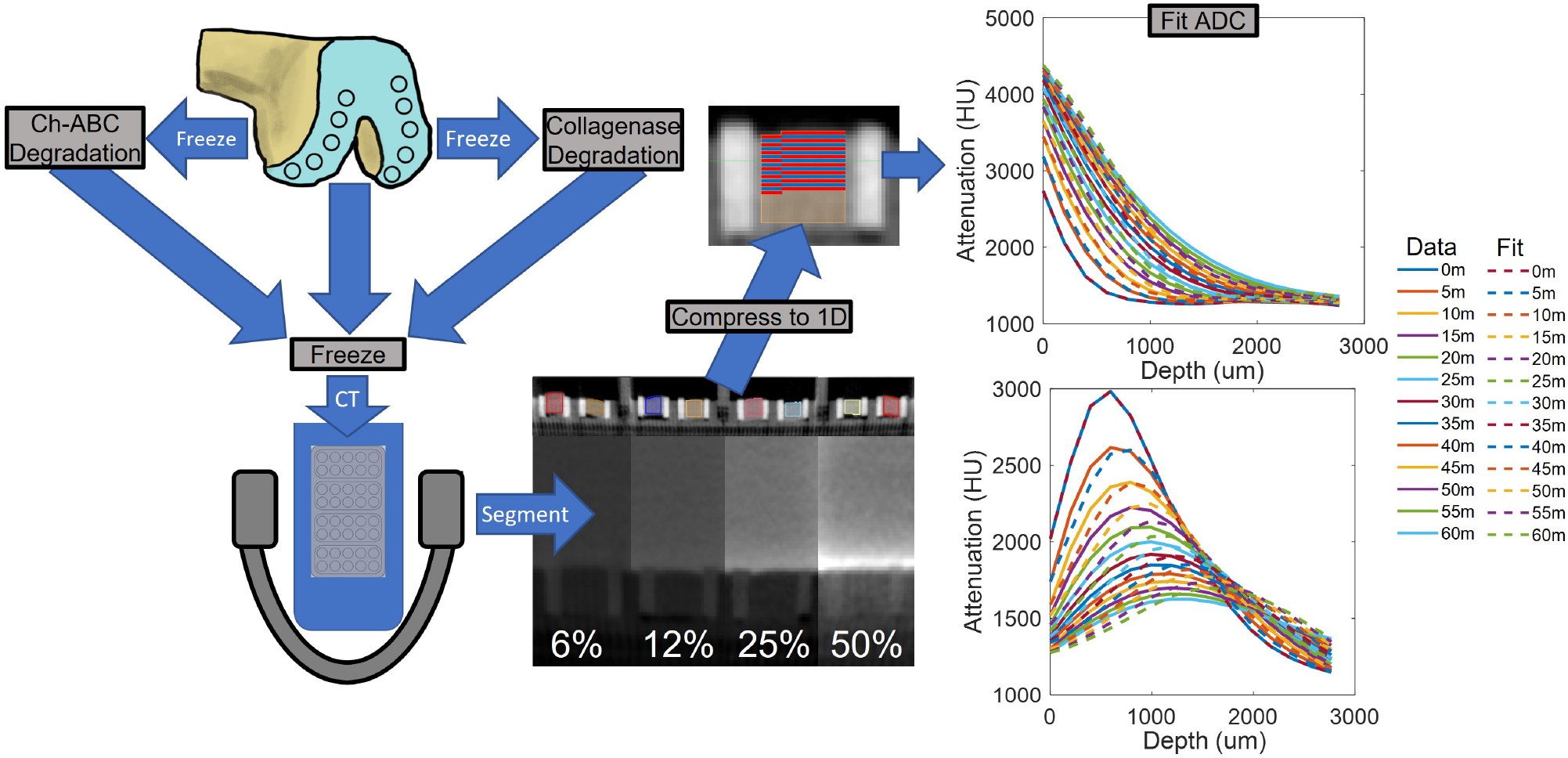
Experimental methods. Explants were extracted from 15 bovine femurs and randomly distributed to untreated, chondroitinase digestion or collagenase digestion groups (n=40/group). Ten explants from each degradation group were exposed to one of four contrast agent dilutions. 1-D line profiles were extracted from reconstructed datasets by averaging across the width of the explant based on distance from the surface. Apparent diffusivity was fit to the line profile data using a least squares minimization routine in MATLAB.

### Materials

Juvenile bovine stifles were from San Jose Valley Veal and Beef (Santa Clara, CA). Protease Inhibitor Cocktail Set I and phosphate buffered saline (PBS) tablets (140mM NaCl, 10mM phosphate buffer, 3mM KCl reconstituted in 1L of deionized water) were from Calbiochem (San Diego, CA). Collagenase 2 and collagenase 4 were from Worthington Biochemical Corporation (Lakewood, NJ). Chondroitinase ABC was from Sigma-Aldrich (St. Louis, MO). Biopsy punches were from Integra Miltex (York, PA). Omnipaque 350 was from GE Healthcare (Chicago, IL).

### Explant Preparation

Full thickness, 5 mm diameter cartilage explants were isolated with a biopsy punch from the medial and lateral femoral condyles (5 explants each) of 15 immature bovine stifle joints and trimmed to a height of 5mm from the articular surface. Explants were frozen in PBS with protease inhibitors and 120 explants were randomly distributed across 3 groups (n=40 each): untreated, chondroitinase ABC digested or collagenase digested. The remaining 30 explants were used in a separate study.

Explants to be digested were thawed at room temperature, transferred to individual wells of a 48-well plate and covered with one of two digest solutions: 0.5mL of PBS with 0.5U/mL chondroitinase ABC or 1mL of PBS with 0.2mg/mL each of collagenase 2 and collagenase 4. Explants were refrigerated at 4°C for 24 hours (chondroitinase) or 30 hours (collagenase) to allow the enzymes to diffuse into the cartilage while inactivate. The plates were then transferred to a water bath and incubated at 25°C for 48 hours (chondroitinase) or 12 hours (collagenase) of digestion. Remaining enzyme was removed by rinsing explants with PBS and equilibration for 24 hours in PBS at 4°C in a new 48-well plate, followed by a second rinse and another 24 hour equilibration. The explants were then individually frozen in PBS plus protease inhibitors until use.

### Explant Scanning

Contrast solutions were prepared by diluting the non-ionic contrast agent (CA) Omnipaque 350 (aqueous solution of 755 mg/mL iohexol providing 350 mg/mL iodine, 1.21 mg/mL tromethamine and 0.1 mg/mL edetate calcium disodium) with PBS to concentrations of 50%, 25%, 12% or 6% v/v. These conditions fall within or below the range of dilutions typically injected for CT arthrography, allowing examination of conditions that may occur as iohexol concentration in the joint drops due to effusion and physiologic clearance. The osmolality of each solution was measured in triplicate using a Wescor 5500 vapor pressure osmometer (Table 1). As in clinical procedures where a non-ionic contrast agent is diluted in saline, the ion concentrations increased as the contrast agent concentration decreased.

**Table 1.** Osmolality and ionic strength of experimental solutions. Osmolality is the mean ± standard deviation of triplicate measurements. Ionic strength is based on reported compositions of Omnipaque 350 and PBS.

| Solution | Omnipaque 350 | PBS | Osmolality (mOsm/kg) | Ionic Strength (mM/L) |
| --- | --- | --- | --- | --- |
| 50% CA | 50% | 50% | 537±3.4 | 83 |
| 25% CA | 25% | 75% | 438±5.7 | 124 |
| 12% CA | 12% | 88% | 366±2.5 | 146 |
| 6% CA | 6% | 94% | 323±1.7 | 156 |
| PBS | 0% | 100% | 287±1.1 | 166 |

Explants from a given digestion group were scanned simultaneously using a c-arm cone beam computed tomography scanner (Artis Zeego, Siemens Healthineers). After thawing at room temperature, each group of 40 explants was placed in individual wells lined with silicone tubing within a custom, 3D-printed fixture and was secured on the base and sides using cyanoacrylate to ensure that only the cartilage surface was exposed to the contrast solution. The fixture was divided into 4 sections, each of which held 10 explants and 45 mL of contrast medium. Explants were kept hydrated with PBS-soaked wipes until immediately prior to the start of the CT scan protocol.

The fixture containing the explants was first scanned without any liquid in the fixture to acquire a reference image that could be used for accurate segmentation. Next, each fixture section was filled with one of the four contrast solutions, followed immediately by an initial scan and subsequent scans every 5 minutes for 1 hour for a total of 13 scans during the contrast absorption phase. The contrast agent bath was then emptied out of the fixture, explant surfaces were blotted dry and a second reference scan was taken with no liquid in the fixture. Each compartment of the fixture was then filled with 45ml PBS and scanned immediately, then scanned every 5 minutes for 1 hour for a total of 13 scans during the contrast desorption phase. The scan protocol was repeated for the intact, chondroitinase-digested and collagenase-digested explants. All scans were acquired at 80kVp using a 20-second protocol with 496 projections. Datasets were reconstructed using the scanner’s included InSpace reconstruction software with isotropic 0.2mm voxels. Each explant was individually segmented from the air scans taken prior to absorption and desorption of the contrast agent using Seg3D2 (version 2.4.3, University of Utah). Averaged 1-D attenuation line profiles for each scan were calculated in Python 3 by averaging attenuation values across the width of the explant for all voxels at a given distance from the segmented cartilage surface, which compensated for the surface of the explants not always being in perfect alignment with the coordinate system of the images.

The apparent diffusivity of each sample was estimated separately for absorption and desorption phases via least squares minimization using the `fmincon` function in MATLAB R2018b (Mathworks, Natick, MA). Diffusivity values were constrained to be between 0 and 10,000*µ*m^2^/s. Homogeneous 1-D Fickian diffusion (Equation 1) was simulated over a 2.5mm depth using the `pdepe` function with 10µm spatial resolution and output time points matching the experimental data:

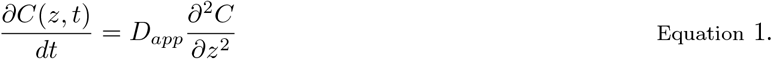

where *C*(*z, t*) is the measured attenuation at a given depth *z* and time *t* and *D*_*app*_ is the apparent diffusivity. Initial conditions were interpolated from the spatial attenuation profile of the first absorption or desorption scan, and time-dependent boundary conditions at the surface (z=0) and lower boundary (z=2.5mm) were interpolated from measured attenuations at those locations. As a goodness of fit measure, the normalized root-mean-square error (nRMSE) for each sample was determined by dividing the RMSE for the absorption or desorption fit by the average attenuation over both phases.

### Data Analysis

One sample (untreated, 50% iohexol group) with an unusually high nRMSE was eliminated after examination of attenuation profiles revealed anomalous patterns due to an unknown artifact. Statistical analyses were performed using Minitab (v20.2, Minitab LLC, State College, PA). The apparent diffusivity and nRMSE were analyzed using repeated measures ANOVA with contrast solution (Omnipaque concentration) and degradation treatment as between-sample factors and diffusion mode (absorption or desorption) as a within-sample factor. Bonferroni’s test was used for pairwise comparisons. Differences were considered statistically significant at p<0.05, and data are presented as mean *±* 95% CI.

### Computational Methods

Simulations of solute transport were used to explore the influences of tissue heterogeneity and non-diffusive transport due to electro-mechano-chemical coupling on the derived apparent diffusivity and to investigate the influences of spatial and temporal scan resolution under idealized conditions in the absence of other experimental artifacts. Figure 2 presents a flowchart detailing the steps in each simulation study.

**Figure 2.**
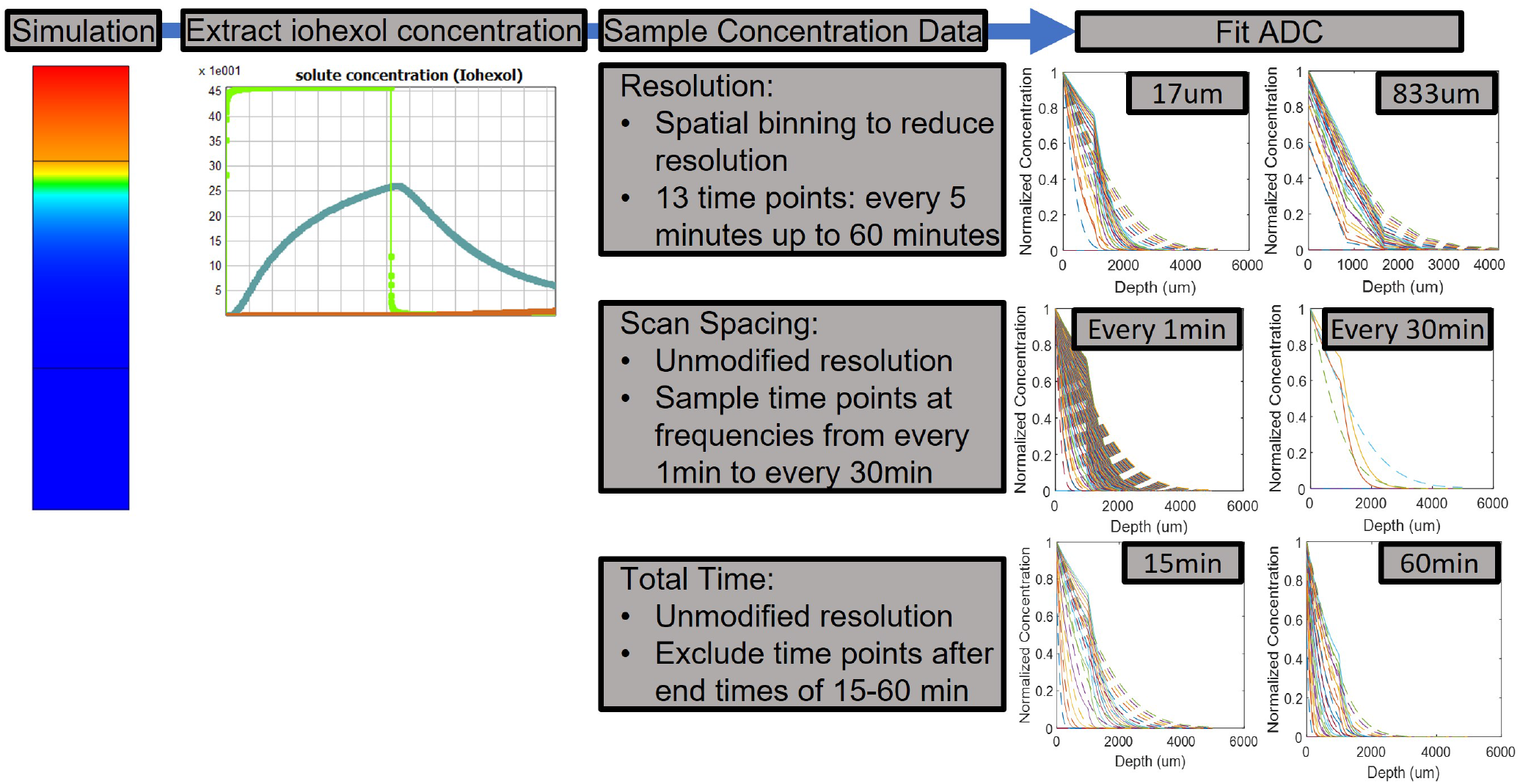
Simulation methods. Simulations were run using FEBio software. Data from the simulation was imported into MATLAB and sampled to reduce data resolution, increase time between simulated scans, or decrease total amount of contrast exposure time.

### Transport Simulations

Finite element models of a cartilage explant were created and solved in FEBio Studio (version 1.5.0, University of Utah)^40^. Four types of simulations were performed: a zonal (3-zone) multiphasic simulation, a homogeneous (1-zone) multiphasic simulation, a zonal diffusion-only simulation, and a homogeneous diffusion-only simulation. Each model consisted of 300 equally sized linear brick elements along the depth of the cartilage, and 1 element along the width of the cartilage.

Cartilage was modeled as a multiphasic, porous material with a fixed negative charge, an ellipsoidal fiber distribution, a neo-Hookean solid matrix, and isotropic, strain-independent permeability and solute diffusivities. To approximate physiologic variation in composition and physical properties, zonal models had three distinct zones with the 20% of the cartilage closest to the surface as the surface layer, the middle 50% as the middle layer, and the final 30% as the deep layer. Homogeneous (1-zone) models used surface zone properties for the full thickness. Cartilage properties for each layer are detailed in Table 2. The model geometry was 0.1 mm x 0.1 mm x 5 mm, with constraints on the sides limiting translation and transport to occur only in the depth direction. Displacement, fluid flow and solute flux were constrained to zero at the model base and unconstrained at the model surface. Bath concentrations of neutral iohexol, Na^+^, and Cl^−^ were prescribed as boundary conditions at the cartilage surface to simulate well-mixed, infinite bath conditions based on the four experimental solutions, as was the corresponding value of the apparent pressure (based on zero fluid pressure in the bath). As concentrations of iohexol and ions simultaneously vary with dilution of the contrast agent in saline, concentrations of iohexol and ions were also manipulated independently and results from that study are presented in the supplement.

**Table 2.** Material properties used for multiphasic modeling of solute transport through cartilage explants

| Property | Zonal Model |  |  | Homogeneous Model |
| --- | --- | --- | --- | --- |
|  | Surface | Middle | Deep |  |
| Solid Volume Fraction (post-swelling) | 0.15 | 0.25 | 0.35 | 0.15 |
| Density ($\text{kg/m}^3$) | 1051 | 1087 | 1123 | 1051 |
| Fixed Charge Density ($\text{mEq/L}$)<sup>41</sup> | -150 | -200 | -250 | -150 |
| Young's Modulus ($\text{kPa}$)<sup>42</sup> | 200 | 500 | 1000 | 200 |
| Poisson's Ratio | 0.1 | 0.1 | 0.1 | 0.1 |
| $\beta_1$<sup>43</sup> | 2.5 | 3.5 | 4.5 | 2.5 |
| $\beta_2$<sup>43</sup> | 2.5 | 3.5 | 4.5 | 2.5 |
| $\beta_3$<sup>43</sup> | 2.5 | 3.5 | 4.5 | 2.5 |
| $\xi_1$ ($\text{MPa}$)<sup>43</sup> | 2.8 | 4.0 | 8.0 | 2.8 |
| $\xi_2$ ($\text{MPa}$)<sup>43</sup> | 1.6 | 2.0 | 4.0 | 1.6 |
| $\xi_3$ ($\text{MPa}$)<sup>43</sup> | 1.6 | 2.0 | 4.0 | 1.6 |
| Permeability ($\text{m}^2/(\text{Pa}\cdot\text{s})$)<sup>44</sup> | 4.55e-15 | 1.46e-15 | 5e-16 | 4.55e-15 |
| Iohexol Diffusivity ($\text{um}^2/\text{s}$) | 228 | 40 | 26 | 228 |
| $\text{Na}^+$ Ion Diffusivity ($\text{um}^2/\text{s}$) | 950 | 842 | 831 | 950 |
| $\text{Cl}^-$ Ion Diffusivity ($\text{um}^2/\text{s}$) | 741 | 618 | 604 | 741 |

### Property Zonal Model Homogeneous Model Surface Middle

The multiphasic simulations consisted of three steps. First, the model was brought to an initial state of equilibrium with an external saline solution. To compensate for swelling effects, the model height was scaled and volume fractions were adjusted so that volume fractions and layer sizes would be equal to those in Table 2 at the end of equilibration. The model started with interstitial fluid and bath concentrations of 154mM NaCl and zero fixed charge density in the cartilage. The fixed charge densities were brought up to the values in Table 2 through a series of intermediate static solutions, at which point the cartilage was at its equilibrium swelling state. Next, boundary concentrations of iohexol, Na+, and Cl- and the corresponding fluid pressure at the surface were changed to values representing one of the four contrast dilutions (Table 3) and 1 hour of contrast agent absorption was simulated. Finally, the iohexol concentration at the surface was reduced to zero while the Na+ and Cl-concentrations were increased to 154mM and a final 1 hour of contrast agent desorption was simulated. During the absorption and desorption phases, the maximum allowed time step size was 15 seconds.

**Table 3.** Boundary concentrations prescribed at the cartilage surface for each step in the simulations

| Step | Na+ (mM) | Cl- (mM) | Iohexol (mM) |
| --- | --- | --- | --- |
| Equilibration | 154 | 154 | 0 |
| Absorption: 50% Iohexol | 77 | 77 | 460 |
| Absorption: 25% Iohexol | 115.5 | 115.5 | 230 |
| Absorption: 12% Iohexol | 135.5 | 135.5 | 115 |
| Absorption: 6% Iohexol | 144.8 | 144.8 | 57.5 |
| Desorption | 154 | 154 | 0 |

The diffusion-only simulations used constrained versions of the corresponding multiphasic models. To eliminate osmotic effects from these simulations, the displacements of all nodes were fixed, the fixed charge density and osmotic coefficients in the tissue were set to zero, the fluid pressure at the cartilage surface was set to zero, and Na+ and Cl-ions were removed from the models. This fully eliminated swelling, so the initial equilibration step and dimensional scaling were unnecessary. As with the multiphasic models, 60 minutes of iohexol absorption was simulated, followed by 60 minutes of iohexol desorption.

### Post-processing

To determine the volume-averaged concentration (which produces the attenuation measured experimentally), interstitial iohexol concentrations were multiplied pointwise by the fluid volume fraction. To simulate variations in CT scan parameters, three different types of post-processing were performed on the zonal multiphasic simulations to produce pseudo-data for analysis in Matlab:

- To independently explore effects of reduced image resolution, iohexol concentrations at 5 minute intervals were spatially binned and averaged with bin sizes ranging from 16.67 to 833.33*µ*m.
- To independently explore effects of sampling frequency, simulation results except for those at specific times were selectively excluded to mimic scan frequencies ranging from once per minute to once every 30 minutes, using iohexol concentrations at all spatial points.
- To independently explore effects of the total diffusion time, the time period covered by the analysis was decreased by selectively excluding simulation results from the absorption and desorption phases after end-times ranging from 15 to 60 minutes.

Each of the zonal multiphasic simulations thus generated 74 pseudo-data sets for both absorption and desorption. For each set of simulated pseudo-data, apparent diffusivities were fit to the simulated attenuation profiles for both absorption and desorption phases. Except for including the full simulation depth and prescribing zero flux boundary conditions at the base, the fit procedure was identical to that described earlier for experimental data.

## RESULTS

### Experimental

Examples of individual explants exposed to different CA concentrations and representative experimental and bestfit line profiles can be found in Figure 1.

Averaged across conditions, the apparent diffusivity of iohexol in untreated cartilage (254.2 *±* 7.2 *µm*^2^*/s*) was consistent with reported values for similarly sized contrast agents^33, 45, 46^. Contrast agent concentration, transport mode (absorption vs. desorption) and tissue digestion all influenced the apparent diffusivity derived from attenuation curves (Fig. 3), with significant interactions between degradation and transport mode and between concentration and transport mode. The interaction between concentration and digestion treatment was not significant, indicating that the relative effects of contrast agent dilution did not significantly vary across digestion groups (and vice-versa).

**Figure 3.**
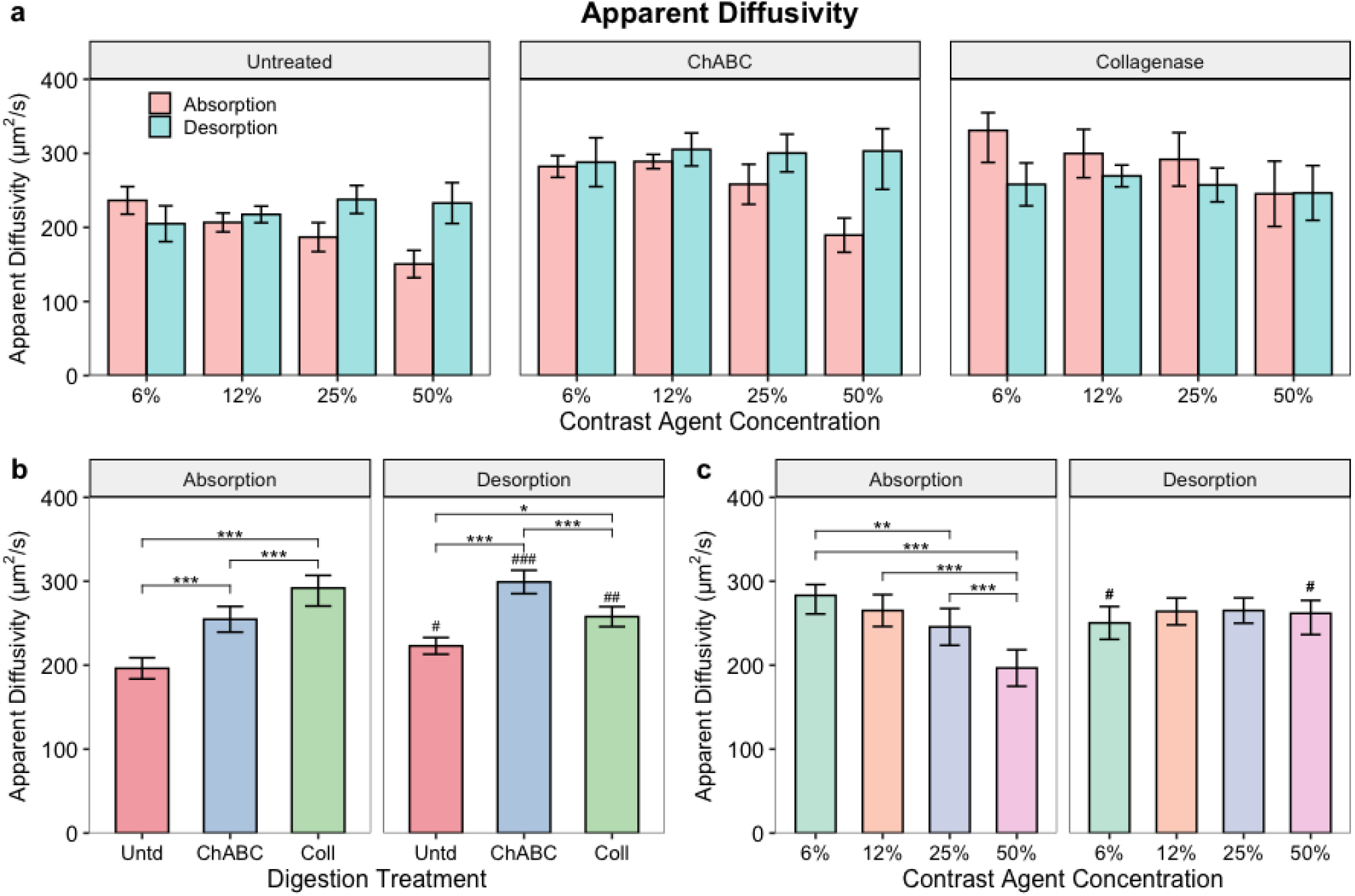
Apparent Diffusivity derived from experimental measurements via least-squares fit. a) Mean values in absorption and desorption modes for each digestion and contrast agent concentration group. Significant interactions were found b) between digestion treatment (Untd = untreated, ChABC = chondroitinase ABC, Coll = collagenase) and transport mode, pooled across contrast agent concentrations and c) between contrast agent concentration and transport mode, pooled across digestion treatments. # p<0.05, ## p<0.01, ### p<0.001 vs. the corresponding absorption group. * p<0.05, ** p<0.01, *** p<0.001 between concentrations or digestion treatment within a transport mode. Data are presented as mean *±* 95% CI.

The apparent diffusivity of chondroitinase- and collagenase-treated explants was significantly higher than that of untreated explants for both absorption and desorption (Figure 3b). Collagenase-treated explants had a higher apparent diffusivity in absorption but a lower apparent diffusivity in desorption compared to chondroitinase-treated explants. For untreated and chondroitinase-treated explants, the apparent diffusivity was greater in desorption than in absorption, while for collagenase-treated explants the apparent diffusivity was lower in desorption than in absorption.

In absorption, the apparent diffusivity decreased with increasing iohexol concentration (Figure 3c), consistent with increased convective retardation of transport due to fluid exudation at higher osmolalities. Apparent diffusivity in absorption was significantly greater at 6% CA than at 25% CA or 50% CA, and was significantly lower at 50% CA than at all other concentrations. The apparent diffusivity in absorption did not significantly vary among concentrations. Consequently, at 6% CA the apparent diffusivity was greater in absorption than in desorption while at 50% CA the apparent diffusivity was greater in desorption than in absorption.

The RMS error normalized by the mean attenuation on a sample-by-sample basis (Figure 4) provides a relative metric of how well the homogeneous Fickian diffusion model describes the transient changes in attenuation. In absorption, the normalized RMS error was significant lower for both enzymatic treatments than for the untreated explants (Figure 4b), indicating less deviation from homogeneous diffusion. This pattern was not maintained in desorption, however, with the chondroitinase-treated explants having significantly greater normalized RMS error than the other conditions. There was a clear, monotonic increase in normalized RMS error with increasing contrast agent concentration (Figure 4c), with significant differences between all concentrations. Consistent with patterns in the apparent diffusivity values, this indicates an increasing deviation from pure diffusive transport with increasing contrast agent concentration and the associated increased bath osmolality and decreased ionic strength.

**Figure 4.**
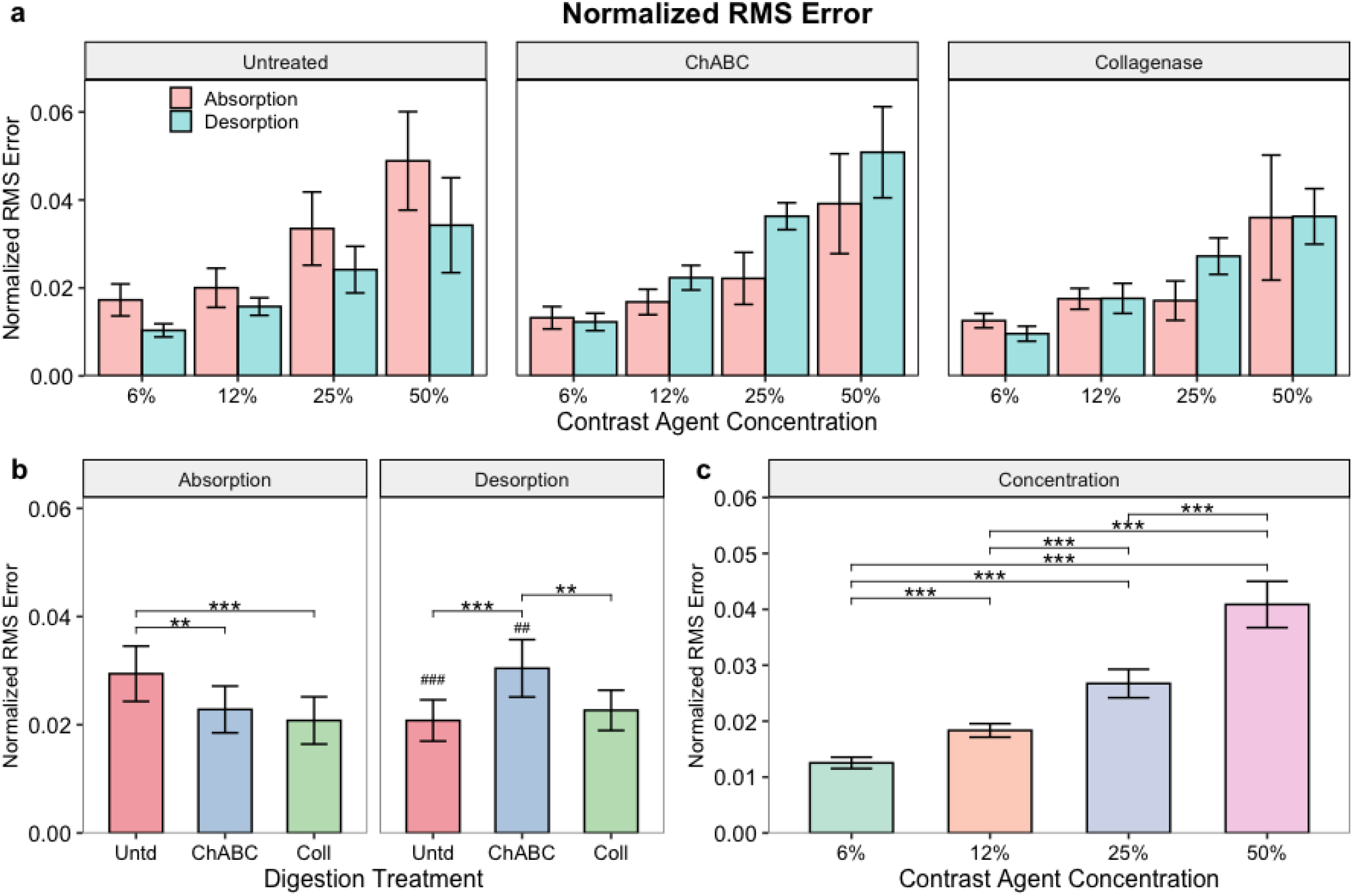
Normalized RMS error for diffusivity fits. a) Mean values in absorption and desorption modes for each digestion and contrast agent concentration group. A significant interaction was found b) between digestion treatment (Untd = untreated, ChABC = chondroitinase ABC, Coll = collagenase) and transport mode, pooled across contrast agent concentrations whereas c) contrast agent concentration had a significant effect independent of transport mode. # p<0.05, ## p<0.01, ### p<0.001 vs. the corresponding absorption group. * p<0.05, ** p<0.01, *** p<0.001 between concentrations or digestion treatments. Data are presented as mean *±* 95% CI.

### Simulations

Examination of diffusivities derived from the different finite element simulations (Figure 5) provides some insights into the mechanisms underlying patterns in the experimental results. Neither the homogeneous nor zonal diffusion-only models exhibited any influence of concentration on the derived diffusivity in either absorption or desorption. In contrast, the homogeneous multiphasic model exhibited small decreases in derived diffusivity with increasing concentration in the absorption phase and the zonal multiphasic model exhibited more pronounced decreases in absorption, while neither exhibited concentration effects on the derived diffusivity in desorption. While different in magnitude, these trends for the multiphasic models are consistent with patterns in the experimental results (Figure 3).

**Figure 5.**
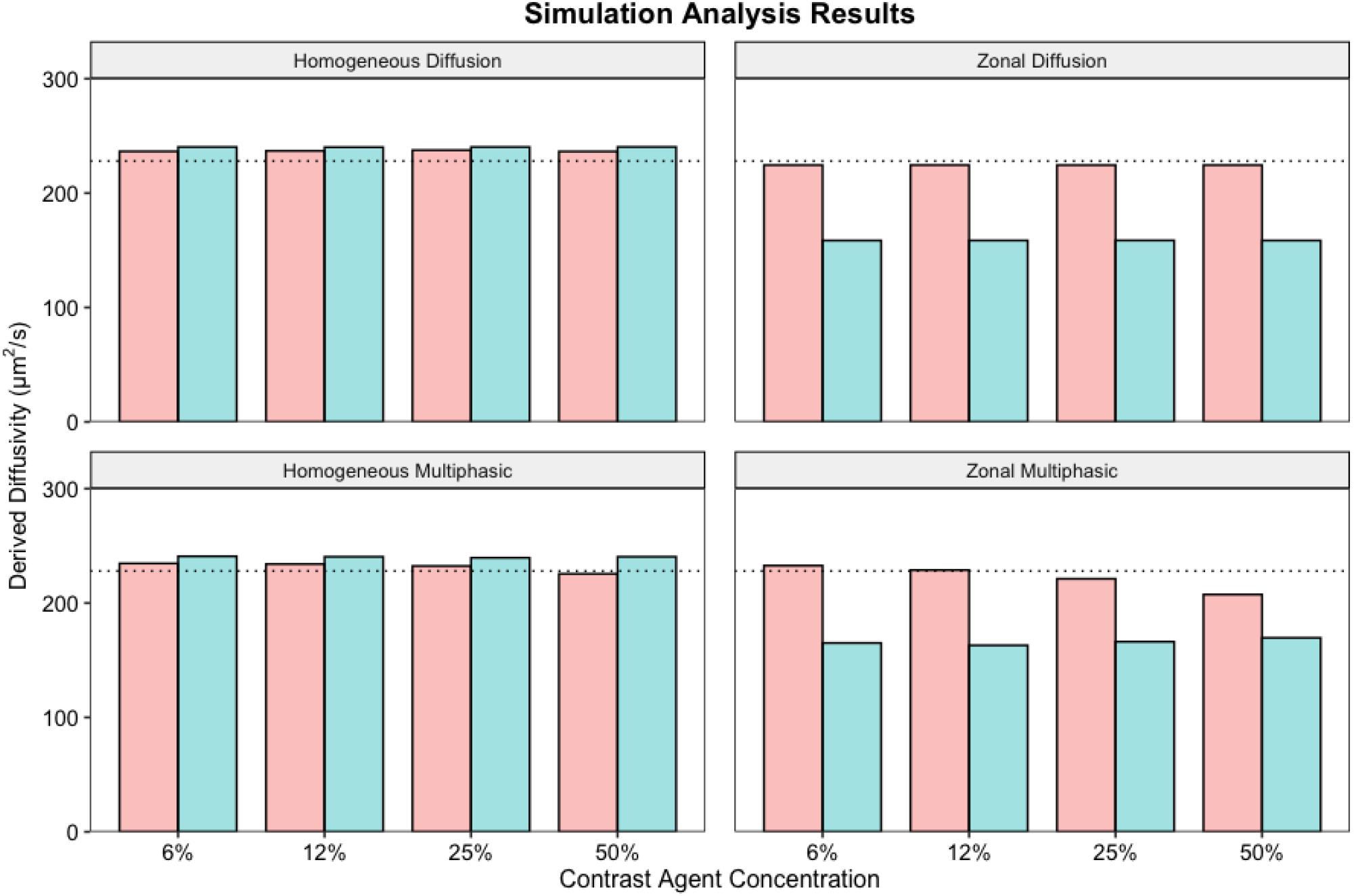
Effects of contrast agent dilution on the diffusivity derived by least-squares fit to predicted attenuation patterns from homogeneous (left) and zonal (right), diffusion-only (top) and multiphasic (bottom) finite element simulations. Dotted lines indicate the iohexol diffusivity assigned to homogeneous models and the surface zone of zonal models.

Neither the homogeneous diffusion-only model nor the homogeneous multiphasic model predicted substantial differences in the derived diffusivity between absorption and desorption phases. In contrast, both of the zonal models produced lower derived diffusivities in desorption than in absorption. As the desorption phase involved both iohexol transport out of the tissue and, especially initially, continued transport of iohexol deeper into the tissue, the lower diffusivity in the middle zone (Table 2) had a greater influence on solute redistribution during the desorption phase than during the preceding absorption phase. Although different from the specific patterns of experimental results, this result suggests that tissue heterogeneity contributes to differences between absorption and desorption in the apparent diffusivity.

The postprocessing variations that simulated aspects of the imaging protocol indicated substantial influences on the absorption diffusivity derived from the zonal multiphasic model. When a spatial resolution decrease was simulated (Figure 6a), the derived diffusivity decreased with increasing voxel size. Conversely, when a temporal resolution decrease was simulated (Figure 6b), the derived diffusivity increased with increasing simulated scan interval (decreasing scan frequency). Finally, simulating a shorter experimental protocol by truncating the simulations at earlier time points (Figure 6c) resulted in higher derived diffusivities, likely because the lower diffusivity in the middle zone has less of an influence on the concentration patterns at earlier times. While the specific interplay among these three factors depends on the actual physical properties of the tissue and the patterns of spatial heterogeneity, these simulation results demonstrate that variations in scan protocol can strongly influence diffusivities derived from CTa. Desorption results, as well as comparable results for the other three models, are presented in the supplement.

**Figure 6.**
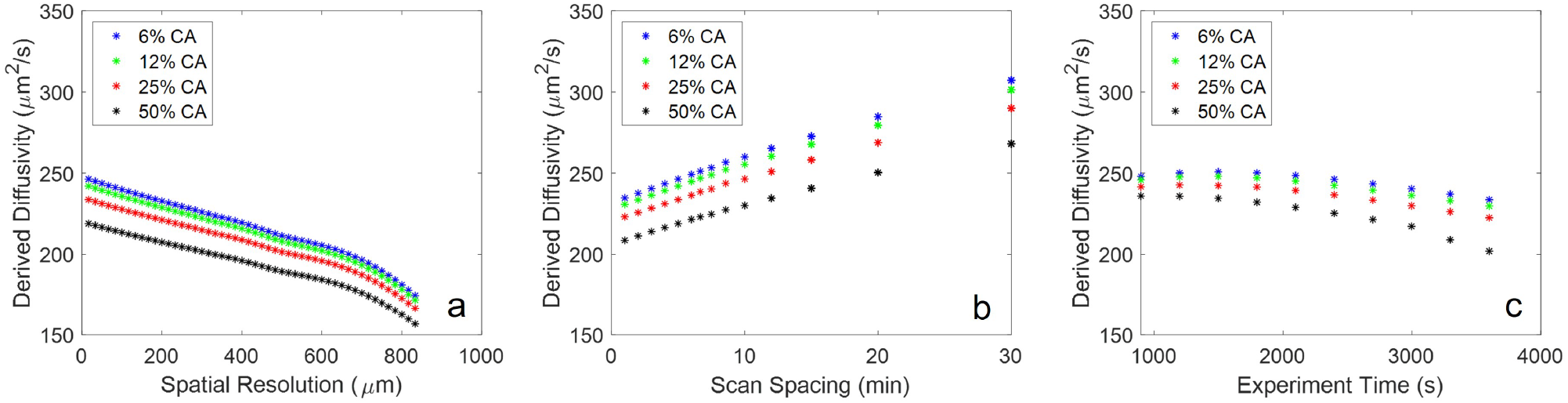
Variations in the apparent diffusivity derived from zonal multiphasic simulations due to simulated changes in (a) spatial scan resolution, (b) interval between sequential scans and (c) total experiment time.

## DISCUSSION

In clinical arthrograms, imaging relatively soon after contrast injection is desirable as the image quality and diagnostic utility progressively deteriorate due both to contrast dilution and blurring of the tissue-fluid interface resulting from contrast uptake by the soft tissues^16, 47^. Early radiographic studies indicated increased contrast uptake by damaged cartilage^47^, and our study supports other recent findings that quantitative analysis of differences in CT-measured contrast agent absorption by cartilage can detect degeneration^28–31, 34, 48^. It also, however, suggests that apparent diffusivity is more sensitive to degeneration than in some previous work^30, 34^ using contrast agents. This may be due in part to differences in the approach to deriving the apparent diffusivity from attenuation data. While many studies fit predictions of the Fickian diffusion model to the bulk average attenuation^30, 33, 35–37^, we took an alternate approach of fitting the model to depth-dependent attenuation profiles, incorporating the measured initial profile as a spatially varying initial condition and using the measured boundary attenuation to specify time-varying boundary conditions. Other studies incorporating measured depth-dependence^49, 50^ have also found increased solute diffusivities in proteoglycan depleted and osteoarthritic cartilage. As in previous use of this approach to derive solute diffusivities from fluorescence recovery after photobleaching (FRAP) data^51^, this reduces the dependence of predicted solute behavior on deviations from ideal theoretical behaviors. In the context of these experimental studies, the line profile approach has the additional advantage of being suitable for analysis of both absorption and desorption, which also may be an issue during in vivo scanning when the intrasynovial solute concentration changes during the scan protocol.

There are some limitations of the controlled model system used in our experiments and differences from conditions present in clinical arthrography scans. We used normal and enzymatically digested juvenile bovine cartilage, which differs in composition and physical properties from those of adolescent or adult human cartilage. While this may lead to some quantitative differences in solute diffusivity and transport kinetics, the basic patterns of solute diffusion in cartilage are conserved across species, with solute size (or mass) being a dominant factor influencing the diffusivity^46^. The primary goal of the two enzymatic treatment protocols was to produce conditions where increased diffusivity was expected, and the apparent diffusivity was significantly greater in both treatment groups than in untreated tissue. Based on a recent characterization^52^, the chondroitinase ABC treatment was expected to remove approximately 50% of the sulfated glycosaminoglycans without directly affecting the collagen network. Collagenase digestion directly degrades the collagen network and results in indirect release of proteoglycans. Although we did not directly assess the degree of degradation produced by our collagenase protocol, the tissue samples from both enzymatic groups differed visibly and palpably from untreated tissue, yet maintained overall integrity. Neither of these treatments replicates the type or pattern of degeneration associated with early-stage osteoarthritis, and studies on human tissues will be required to determine how well this approach translates to detection of clinically realistic degradation patterns.

Our results demonstrate that variations in Omnipaque 350 dilution with saline produced substantial variations in the apparent diffusivity derived from attenuation profiles. For untreated tissues, the apparent diffusivity in absorption for the 50% CA group was 36% lower than that of the most dilute (and closest to ideal conditions) 6% CA group. For context, at 6% CA the apparent diffusivities of chondroitinase ABC- and collagenase-treated explants were 24% and 44% higher, respectively, than that of untreated tissue. Similar concentration trends were seen in the multiphasic finite element simulations but not in the diffusion-only simulations, supporting the role of osmotically-induced fluid flow in reducing early solute transport into the tissue and thus decreasing the apparent diffusivity when fit with the Fickian model. This is also consistent with recent observations that these solutions can rapidly alter the mechanical behaviors of cartilage and meniscus^52, 53^, also due in large part to osmotic effects. The particular contrast agent chosen and preparation approach will have a large influence on the influence of the contrast solution on diffusivity measurement. For example, previous studies involving anionic contrast agents^35^ did not find a relationship between concentration and apparent diffusivity in solutions adjusted to be isoosmolar. Contrast solutions for CT arthrography are based on a variety of ionic and nonionic contrast agents and span a wide range in both osmolality and ionic strength^53^, and many of these solutions would be expected to produce substantial deviations from diffusive transport. The choice of contrast agent for CTa diffusion measurement may depend largely on practical considerations (e.g., institutional preference, regulatory approval or reluctance to use ionic contrast agents in the United States), and the ability to minimize or quantify the degree of non-Fickian behaviors for a given contrast agent will influence the ability to accurately detect degenerative changes. Previous experimental studies indicated that diffusivity in absorption and desorption are similar^45, 54^, except for solutes that bind to the cartilage matrix^32, 55^. In this study, differences in the apparent diffusivity of iohexol between absorption and desorption are likely primarily due to osmotic effects at higher concentrations. Most of this difference is due to effects during absorption, as the large initial difference in osmolality between the bath and the tissue interstitial fluid drives fluid flow out of the tissue. Despite the iohexol uptake and ion flows that had occurred, the osmotic difference between the tissue interior and the PBS bath at the start of desorption were relatively small, with the result that apparent diffusivities during the desorption phase were fairly insensitive to the previous CA concentration. The iodine content of the contrast solution also influences the extent to which imaging artifacts affect the ability to characterize solute diffusion. Higher iodine concentrations, such as in the 50% CA solution used in this work, can cause beam hardening^23, 28, 36, 56^ and associated errors in the measured attenuation. On the other hand, low iodine concentrations could result in a lower effective signal-to-noise ratio and lead to difficulties in distinguishing healthy and unhealthy cartilage, in addition to poorer diagnostic quality for morphology evaluation. In practice, the contrast agent concentration in the synovial space will vary with time due to effusion and physiologic clearance, both of which are likely to vary patient-to-patient. Exposure of explants to a dynamic boundary condition where contrast agent concentration at the cartilage surface decreases over time as it would in vivo^57–59^ could give additional information about how robust apparent diffusivity is as a biomarker for degradation detection. The emergence of new CT technologies such as photon counting detectors have the potential to both reduce beam hardening artifacts and allow accurate measurements at lower attenuation, both of which would be beneficial for diffusion measurement.

The multiphasic finite element models provide qualitative insights into the phenomena responsible for experimental findings, but incorporate many simplifications of actual tissue behaviors. For example, neither the hydraulic permeability nor the solute diffusivities are modeled as strain-dependent, while in reality both would decrease substantially with tissue compression. Particularly for the higher concentrations, the initial osmotically driven fluid efflux would be accompanied by tissue compression near the surface, slowing both fluid efflux and contrast uptake and further reducing the apparent diffusivity. Additionally, while the zonal multiphasic model produced behaviors generally consistent with experimental findings, the heterogeneity in biophysical properties such as diffusivity^37, 60, 61^ is both more gradual and more complex than the simple 3-zone representation in these models. As the desorption phase of our experiments involved simultaneous solute transport out of the tissue surface and continued transport into deeper layers, the specific heterogeneity pattern may have a stronger influence on the desorption response. While solute diffusivities are generally lower in the deeper, lower porosity regions of cartilage, some solutes exhibit increased peak diffusivities 200-300µm beneath the surface^62^, which could substantially alter the differences between absorption and desorption. Adding to this complexity, patterns of heterogeneity change with tissue degeneration. It may be possible to develop imaging and analysis strategies that leverage these phenomena, providing insights into tissue status based on differences between responses in different regimes (e.g, early absorption with high concentrations, late desorption with lower concentrations). Additionally, incorporation of depthwise variations (such as piecewise constant or parametric depth dependent diffusivities) may be better suited to fitting experimental data than a simple model with a single apparent diffusivity.

We used the zonal multiphasic model to explore the influence of various CT scan protocol variations on the diffusivity derived using the Fickian diffusion model. Reducing spatial and temporal resolution both affected derived diffusivity, with opposite effects of coarser temporal and spatial resolution. These effects were independent of the CA concentration, heterogeneity, and inclusion of multiphasic phenomena (see supplement). While the specific implications may depend on the exact contrast solution, tissue state and other variables, it is clear that variations in scan parameters themselves can influence the derived diffusivity as much as changes in tissue composition or external conditions. This indicates the importance of consistent experimental protocols within a study or within a given clinical center, and also impact the relationship between laboratory and clinical measurements because imaging systems used in explant studies often provide much higher resolution^50, 63^ than clinical scanners. Any comparisons across scanners would require careful development of calibration protocols, and repeated scans of a given subject would have to follow consistent protocols for reliable longitudinal assessment. The duration of the transport response captured also influenced the derived diffusivity, with apparent convergence as the period became substantially longer than the initial transient. The effects of total diffusion time on the derived diffusivity may also be related to tissue heterogeneity, as longer diffusion times allowed more contrast to reach deeper layers of the cartilage with lower diffusivities, therefore reducing the apparent diffusivity derived with the Fickian model. With relatively few exceptions^24, 25, 27^, explant studies have usually involved near-equilibrium time scales^19, 26, 30, 34–37,39^ which are not feasible in vivo and may mask unusual behaviors at earlier time points (such as the concentration dependence that we found). In vivo protocols on shorter time scales^28, 29, 31^ have typically reported attenuation or partition, but have not derived diffusion coefficients.

This study differs from many previous studies exploring transport of contrast agents in cartilage because it involved clinically relevant timescales when the attenuation change within the tissue is strongly influenced by the surface properties. This is an important step in translating apparent diffusivity measurements from well controlled explant experiments to a quantitative CT biomarker that can be utilized for evaluation of cartilage health in vivo. Both degradation protocols used in this study were relatively mild and involved no physical disruption of the surface, supporting further studies on CTa diffusion measurement as a biomarker for early cartilage degeneration associated with osteoarthritis.

## CONCLUSIONS

This study demonstrates the feasibility of using apparent diffusivity calculated by tracking diffusion of a neutral solute into cartilage as a biomarker for detecting degradation in explants subject to mild degradation protocols. Properties of the contrast solution can substantially influence the apparent diffusivity, and such effects should be minimized or understood well enough to allow analytical compensation. Scan protocol parameters such as spatial and temporal scan resolution and the time period covered by scans can also influence the calculated apparent diffusivity, so it will be important to establish consistent imaging protocols and calibration procedures. With imaging and analysis procedures tailored to minimize the influence of common artifacts, CTa-based diffusivity measurement holds promise as an imaging biomarker for detecting early degenerative changes.

## SUPPLEMENTAL INFORMATION 1: INDIVIDUAL EFFECTS OF ION AND IOHEXOL CONCENTRATION

In the main text of this paper, the effects of exposing cartilage explants to iohexol diluted to varying degrees with saline are investigated. Varying the concentrations of ions and iohexol have different effects that interact, and changing them simultaneously could cause unexpected patterns in apparent diffusivity variation with concentration changes. Simulation in FEBio was used to parse the effects of ion concentration and iohexol concentration on diffusion of iohexol into articular cartilage.

All simulations were performed in FEBio using the zonal multiphasic model described in the main text. In the first set of simulations, ion concentration was held constant at the equivalent of 0.75X saline and iohexol concentration was varied from 0.1X-10X the concentration of in Omnipaque 350. In the second set of simulations, iohexol concentration was held at 25% by volume and concentrations of Na+ and Cl-were varied from the equivalent of 0.1X-10X saline. An apparent diffusivity was fit to the simulation results by fitting the Fickian diffusion equation to the data with a least squares optimization routine in MATLAB (R2018a), as described in the main text. The results of these simulations can be found in Figure 7.

**Figure 7.**
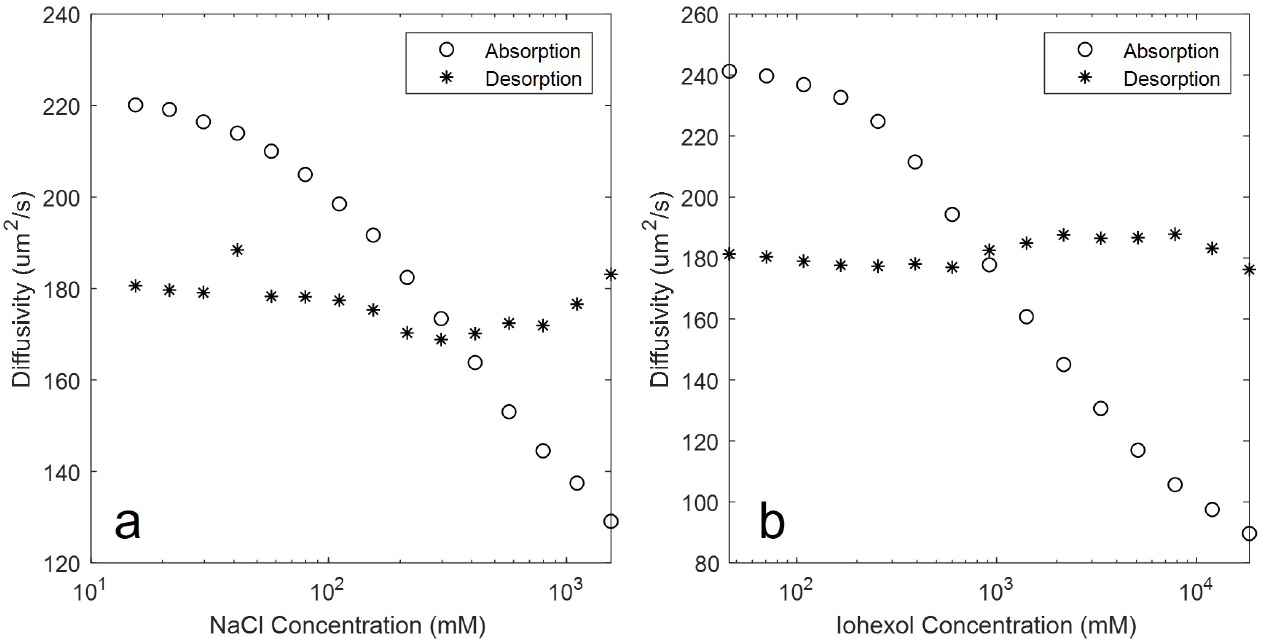
Apparent diffusivity derived from zonal multiphasic finite element simulations varied with (a) ion concentration at fixed iohexol concentration and (b) iohexol concentration for fixed ion concentrations.

Figure 7 shows that the apparent diffusivity derived from zonal multphasic model simulations decreased as either iohexol concentration or ion concentrations increased. This effect is best explained by examining fluid flux at the extreme ends of the concentration spectrum simulated, as shown in Figure 8. Whether due to purely chemical effects (iohexol) or combined electrochemical effects (saline), high bath concentrations induced initial fluid efflux that abated with time, while low bath concentrations induced fluid uptake that enhanced iohexol uptake.

**Figure 8.**
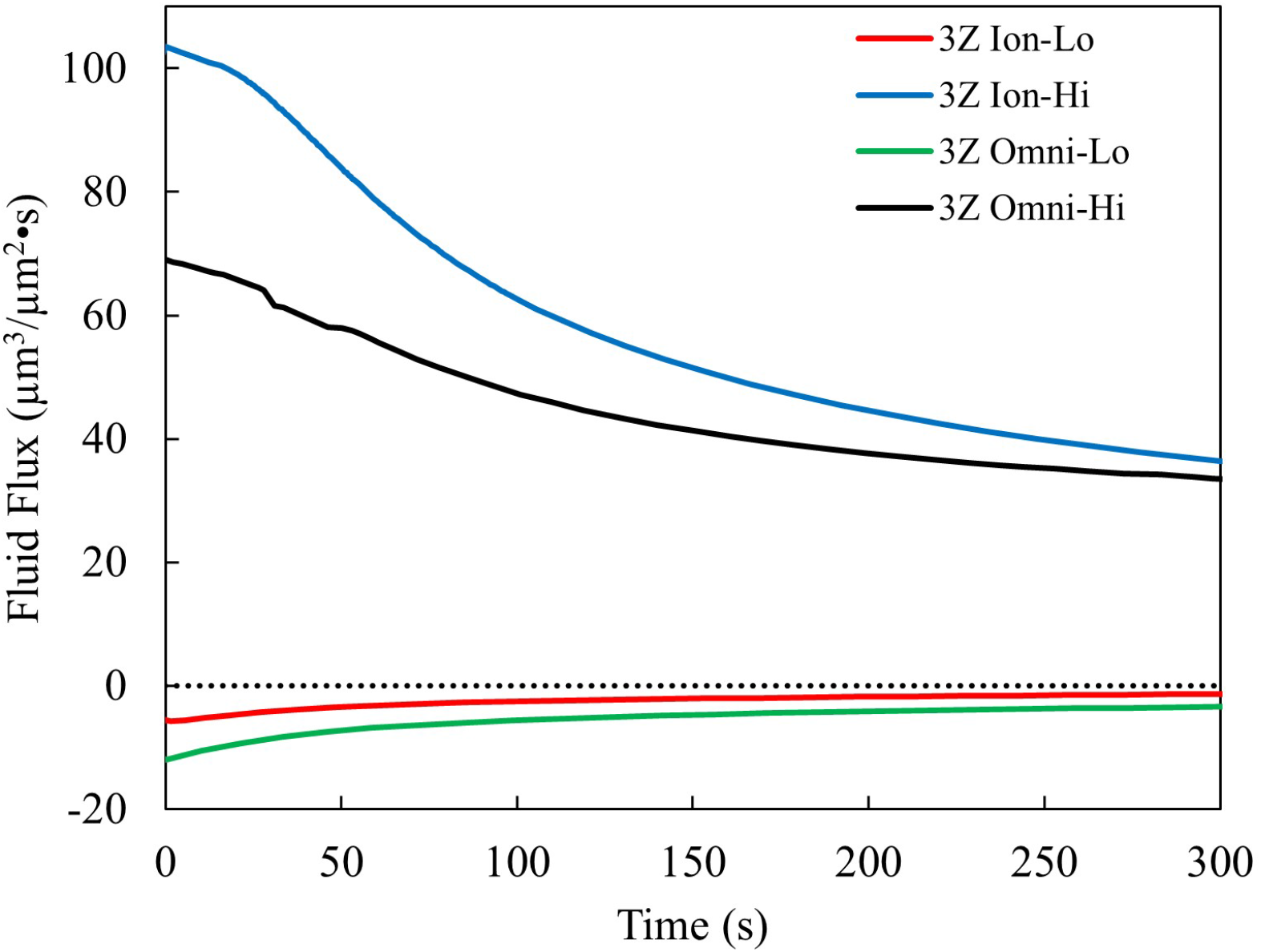
Average fluid flux in the 20% of cartilage closest to the surface for 5 minutes post exposure to solution. Plots are of lowest and highest simulated concentrations of iohexol and NaCl for the zonal multiphasic model. Positive flux denotes flux towards the cartilage surface.

In simulations with higher concentrations of solutes, regardless of whether they were Na+ and Cl-ions or iohexol, fluid flow towards the cartilage surface retards iohexol flux into the cartilage, resulting in lower apparent iohexol diffusivity.

## SUPPLEMENTAL INFORMATION 2: DERIVED DIFFUSIVITY VS. ACQUISITION PARAMETERS IN ALL MODELS

In the main text, we describe effects of simulated variations in scan parameters on the apparent diffusivity derived from the absorption phase of zonal multiphasic model simulations. Here we present effects of these variations on the apparent diffusivity in both absorption and desorption derived from multiphasic and diffusion-only simulations with zonal (Figure 9) and homogeneous (Figure 10) material property distributions.

**Figure 9.**
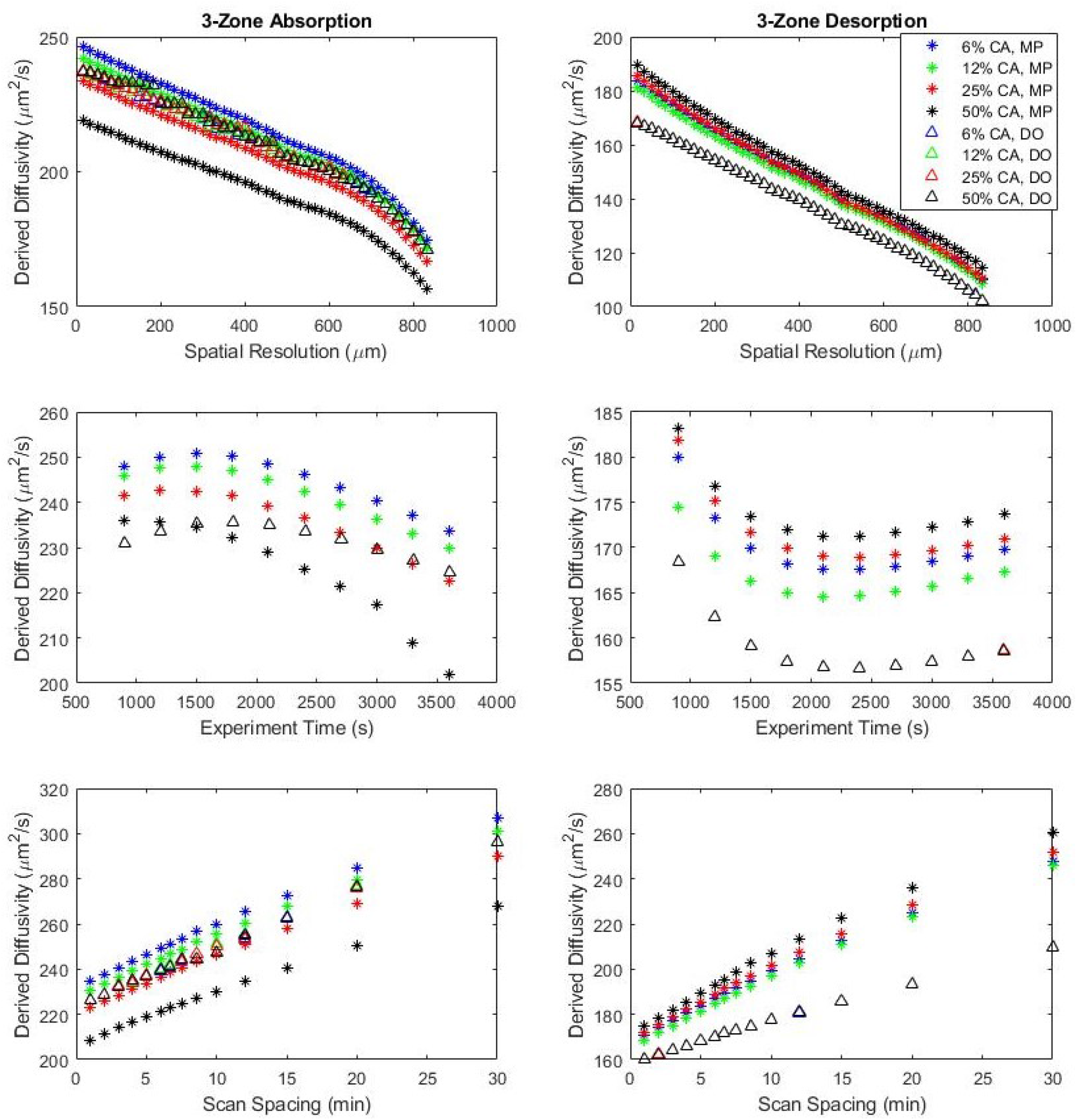
Derived diffusivity variation with spatial resolution, experiment time, and temporal scan spacing for zonal multiphasic (MP) and diffusion-only (DO) finite element models. Note differences in scale for different plots.

**Figure 10.**
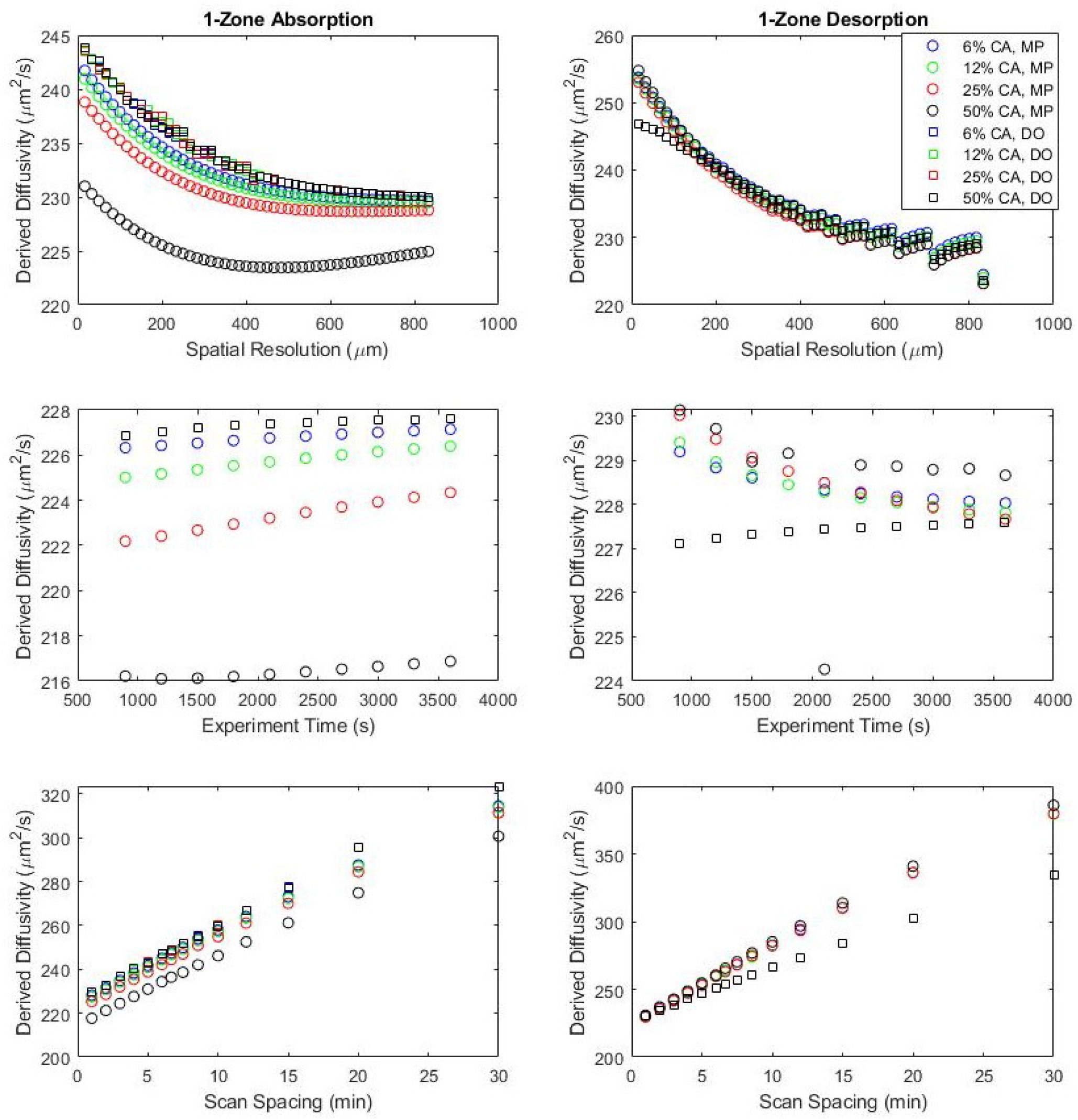
Derived diffusivity variation with spatial resolution, experiment time, and temporal scan spacing for homogeneous multiphasic (MP) and diffusion-only (DO) finite element models. Note differences in scale for different plots

The effects of spatial resolution (voxel size) and temporal resolution (time between scans) were similar across models, absorption and desorption, and CA concentrations. This suggests that relative comparisons among samples, patients, or studies with varied tissue behaviors would be valid if spatial and temporal resolution are consistent. However, comparisons among studies where these parameters are not identical should be approached with caution, as effects of resolution differences could mask (or be misinterpreted as) differences in diffusivity. Varying the total scan period also affected the apparent diffusivity, but the effects were less consistent across models.

## REFERENCES

1. Bitton R. The economic burden of osteoarthritis. The American Journal of Managed Care. 2009;15(8 Suppl):S230–5.

2. Michael JWP, Schlüter-Brust KU, Eysel P. The epidemiology, etiology, diagnosis, and treatment of osteoarthritis of the knee. Deutsches Arzteblatt International. 2010;107(9):152.

3. Fife RS, Brandt KD, Braunstein EM, Katz BP, Shelbourne KD, Kalasinski LA, et al. Relationship between arthroscopic evidence of cartilage damage and radiographic evidence of joint space narrowing in early osteoarthritis of the knee. Arthritis & Rheumatism. 1991;34(4):377–382.

4. Sophia Fox AJ, Bedi A, Rodeo SA. The basic science of articular cartilage: structure, composition, and function. Sports Health. 2009;1(6):461–468.

5. Martel-Pelletier J, Boileau C, Pelletier JP, Roughley PJ. Cartilage in normal and osteoarthritis conditions. Best Practice & Research Clinical Rheumatology. 2008;22(2):351–384.

6. Hjertquist SO, Lemperg R. Identification and concentration of the glycosaminoglycans of human articular cartilage in relation to age and osteoarthritis. Calcified Tissue Research. 1972;10(1):223–237.

7. Guilak F, Ratcliffe A, Lane N, Rosenwasser MP, Mow VC. Mechanical and biochemical changes in the superficial zone of articular cartilage in canine experimental osteoarthritis. Journal of Orthopaedic Research. 1994;12(4):474–484.

8. Dijkgraaf LC, de Bont LG, Boering G, Liem RS. The structure, biochemistry, and metabolism of osteoarthritic cartilage: a review of the literature. Journal of Oral and Maxillofacial Surgery. 1995;53(10):1182–1192.

9. Quere JB, Phan C, Miquel A, Li L, Arrivé L, Menu Y, et al. MDCT arthrography assessment of the severity of cartilage damage and scapholunate dissociation in regard to specific-component tears of the scapholunate interosseous ligament. European Journal of Radiology. 2020 apr;125:108901.

10. Vopat ML, Hermanns CA, Midtgaard KS, Baker J, Coda RG, Cheema SG, et al. Imaging modalities for the glenoid track in recurrent shoulder instability: A systematic review. Orthopaedic Journal of Sports Medicine. 2021 jun;9(6):23259671211006750.

11. Dornberger JE, Rademacher G, Stengel D, Hönning A, Dipl-Phys GS, Eisenschenk A, et al. What Is the Diagnostic Accuracy of Flat-panel Cone-beam CT Arthrography for Diagnosis of Scapholunate Ligament Tears? Clinical Orthopaedics and Related Research. 2021 jan;479(1):151–160.

12. Kim S, Lee GY, Lee JS. Evaluation of the triangular fibrocartilage: comparison of two-compartment wrist CT arthrography using the distal radioulnar and radiocarpal joints and unicompartment wrist CT arthrography using the radiocarpal joint. The British Journal of Radiology. 2019 oct;92(1102):20190298.

13. Lee GY, Kim S, Baek SH, Jang EC, Ha YC. Accuracy of magnetic resonance imaging and computed tomography arthrography in diagnosing acetabular labral tears and chondral lesions. Clinics in Orthopedic Surgery. 2019 mar;11(1):21–27.

14. Lenoir H, Carlier Y, Ferrand M, Vidil A, Desmoineaux P, Society FA. Can preoperative imaging predict the outcomes after arthroscopic release for elbow arthritis? Orthopaedics & Traumatology, Surgery & Research. 2019 ec;105(8S):S229–S234.

15. Pirimoglu B, Ogul H, Polat G, Kantarci M, Levent A. The comparison of direct magnetic resonance arthrography with volumetric interpolated breath-hold examination sequence and multidetector computed tomography arthrography techniques in detection of talar osteochondral lesions. Acta Orthopaedica et Traumatologica Turcica. 2019 may;53(3):209–214.

16. Obermann WR, Kieft GJ. Knee arthrography: a comparison of iohexol, ioxaglate sodium meglumine, and metrizoate. Radiology. 1987 mar;162(3):729–733. Available from: http://dx.doi.org/10.1148/radiology.162.3.3544035.

17. Ingram C, Stoker DJ. Contrast media in double-contrast arthrography of the knee: a comparison of ioxaglate and iothalamate preparations. The British Journal of Radiology. 1986 feb;59(698):143–146.

18. Siebelt M, van Tiel J, Waarsing J, Piscaer T, van Straten M, Booij R, et al. Clinically applied CT arthrography to measure the sulphated glycosaminoglycan content of cartilage. Osteoarthritis and Cartilage. 2011;19(10):1183–1189.

19. Bansal P, Stewart R, Entezari V, Snyder B, Grinstaff M. Contrast agent electrostatic attraction rather than repulsion to glycosaminoglycans affords a greater contrast uptake ratio and improved quantitative CT imaging in cartilage. Osteoarthritis and Cartilage. 2011;19(8):970–976.

20. Joshi NS, Bansal PN, Stewart RC, Snyder BD, Grinstaff MW. Effect of contrast agent charge on visualization of articular cartilage using computed tomography: exploiting electrostatic interactions for improved sensitivity. Journal of the American Chemical Society. 2009;131(37):13234–13235.

21. Lakin BA, Patel H, Holland C, Freedman JD, Shelofsky JS, Snyder BD, et al. Contrast-enhanced CT using a cationic contrast agent enables non-destructive assessment of the biochemical and biomechanical properties of mouse tibial plateau cartilage. Journal of Orthopaedic Research. 2016;34(7):1130–1138.

22. Palmer AW, Guldberg RE, Levenston ME. Analysis of cartilage matrix fixed charge density and threedimensional morphology via contrast-enhanced microcomputed tomography. Proceedings of the National Academy of Sciences. 2006;103(51):19255–19260.

23. Honkanen MK, Saukko AE, Turunen MJ, Shaikh R, Prakash M, Lovric G, et al. Synchrotron microCT reveals the potential of the dual contrast technique for quantitative assessment of human articular cartilage composition. Journal of Orthopaedic Research. 2020;38(3):563–573.

24. Saukko AE, Turunen MJ, Honkanen MK, Lovric G, Tiitu V, Honkanen JT, et al. Simultaneous quantitation of cationic and non-ionic contrast agents in articular cartilage using synchrotron microCT imaging. Scientific Reports. 2019;9(1):1–9.

25. Saukko AE, Honkanen JT, Xu W, Väänänen SP, Jurvelin JS, Lehto VP, et al. Dual contrast CT method enables diagnostics of cartilage injuries and degeneration using a single CT image. Annals of Biomedical Engineering. 2017;45(12):2857–2866.

26. Bhattarai A, Honkanen JT, Myller KA, Prakash M, Korhonen M, Saukko AE, et al. Quantitative dual contrast CT technique for evaluation of articular cartilage properties. Annals of Biomedical Engineering. 2018;46(7):1038–1046.

27. Honkanen MK, Saukko AE, Turunen MJ, Xu W, Lovric G, Honkanen JT, et al. Triple contrast CT method enables simultaneous evaluation of articular cartilage composition and segmentation. Annals of Biomedical Engineering. 2020;48(2):556–567.

28. Kokkonen HT, Aula AS, Kröger H, Suomalainen JS, Lammentausta E, Mervaala E, et al. Delayed computed tomography arthrography of human knee cartilage in vivo. Cartilage. 2012;3(4):334–341.

29. Myller KA, Turunen MJ, Honkanen JT, Väänänen SP, Iivarinen JT, Salo J, et al. In vivo contrast-enhanced cone beam CT provides quantitative information on articular cartilage and subchondral bone. Annals of Biomedical Engineering. 2017;45(3):811–818.

30. Kokkonen H, Jurvelin J, Tiitu V, Töyräs J. Detection of mechanical injury of articular cartilage using contrast enhanced computed tomography. Osteoarthritis and Cartilage. 2011;19(3):295–301.

31. Kokkonen HT, Suomalainen JS, Joukainen A, Kröger H, Sirola J, Jurvelin JS, et al. In vivo diagnostics of human knee cartilage lesions using delayed CBCT arthrography. Journal of Orthopaedic Research. 2014;32(3):403–412.

32. DiDomenico CD, Lintz M, Bonassar LJ. Molecular transport in articular cartilage—what have we learned from the past 50 years? Nature Reviews Rheumatology. 2018;14(7):393–403.

33. Kulmala K, Korhonen R, Julkunen P, Jurvelin J, Quinn T, Kröger H, et al. Diffusion coefficients of articular cartilage for different CT and MRI contrast agents. Medical Engineering & Physics. 2010;32(8):878–882.

34. Kokkonen H, Mäkelä J, Kulmala K, Rieppo L, Jurvelin J, Tiitu V, et al. Computed tomography detects changes in contrast agent diffusion after collagen cross-linking typical to natural aging of articular cartilage. Osteoarthritis and Cartilage. 2011;19(10):1190–1198.

35. Silvast TS, Jurvelin JS, Tiitu V, Quinn TM, Töyräs J. Bath concentration of anionic contrast agents does not affect their diffusion and distribution in articular cartilage in vitro. Cartilage. 2013;4(1):42–51.

36. Arbabi V, Pouran B, Weinans H, Zadpoor AA. Multiphasic modeling of charged solute transport across articular cartilage: Application of multi-zone finite-bath model. Journal of Biomechanics. 2016;49(9):1510–1517.

37. Arbabi V, Pouran B, Weinans H, Zadpoor A. Transport of neutral solute across articular cartilage: the role of zonal diffusivities. Journal of Biomechanical Engineering. 2015;137(7).

38. Saarakkala S, Julkunen P, Kiviranta P, Mäkitalo J, Jurvelin J, Korhonen R. Depth-wise progression of osteoarthritis in human articular cartilage: investigation of composition, structure and biomechanics. Osteoarthritis and Cartilage. 2010;18(1):73–81.

39. Pouran B, Arbabi V, Zadpoor AA, Weinans H. Isolated effects of external bath osmolality, solute concentration, and electrical charge on solute transport across articular cartilage. Medical Engineering & Physics. 2016;38(12):1399–1407.

40. Maas SA, Ellis BJ, Ateshian GA, Weiss JA. FEBio: finite elements for biomechanics. Journal of Biomechanical Engineering. 2012;134(1).

41. Chen S, Falcovitz Y, Schneiderman R, Maroudas A, Sah R. Depth-dependent compressive properties of normal aged human femoral head articular cartilage: relationship to fixed charge density. Osteoarthritis and Cartilage. 2001;9(6):561–569.

42. Schinagl RM, Gurskis D, Chen AC, Sah RL. Depth-dependent confined compression modulus of full-thickness bovine articular cartilage. Journal of Orthopaedic Research. 1997;15(4):499–506.

43. Ateshian GA, Rajan V, Chahine NO, Canal CE, Hung CT. Modeling the Matrix of Articular Cartilage Using a Continuous Fiber Angular Distribution Predicts Many Observed Phenomena. Journal of Biomechanical Engineering. 2009 04;131(6). 061003. Available from: https://doi.org/10.1115/1.3118773.

44. Chen A, Bae W, Schinagl R, Sah R. Depth-and strain-dependent mechanical and electromechanical properties of full-thickness bovine articular cartilage in confined compression. Journal of Biomechanics. 2001;34(1):1–12.

45. Shafieyan Y, Khosravi N, Moeini M, Quinn TM. Diffusion of MRI and CT contrast agents in articular cartilage under static compression. Biophysical Journal. 2014;107(2):485–492.

46. DiDomenico CD, Bonassar LJ. How can 50 years of solute transport data in articular cartilage inform the design of arthritis therapeutics? Osteoarthritis and Cartilage. 2018 nov;26(11):1438–1446. Available from: http://dx.doi.org/10.1016/j.joca.2018.07.006.

47. Roebuck EJ. Double contrast knee arthrography some new points of technique including the use of dimer X. Clinical Radiology. 1977 jan;28(3):247–257. Available from: https://linkinghub.elsevier.com/retrieve/pii/S0009926077801712.

48. Decker SG, Moeini M, Chin HC, Rosenzweig DH, Quinn TM. Adsorption and distribution of fluorescent solutes near the articular surface of mechanically injured cartilage. Biophysical Journal. 2013;105(10):2427–2436.

49. Torzilli PA, Arduino JM, Gregory JD, Bansal M. Effect of proteoglycan removal on solute mobility in articular cartilage. Journal of Biomechanics. 1997;30(9):895–902.

50. Meng H, Quan Q, Yuan X, Zheng Y, Peng J, Guo Q, et al. Diffusion of neutral solutes within human osteoarthritic cartilage: Effect of loading patterns. Journal of Orthopaedic Translation. 2020 may;22:58–66.

51. Irrechukwu ON, Levenston ME. Improved estimation of solute diffusivity through numerical analysis of FRAP experiments. Cellular and Molecular Bioengineering. 2009 mar;2(1):104–117. Available from: http://link.springer.com/10.1007/s12195-009-0042-1.

52. Baylon EG, Levenston ME. Osmotic Swelling Responses Are Conserved Across Cartilaginous Tissues With Varied Sulfated-Glycosaminoglycan Contents. Journal of Orthopaedic Research. 2020;38(4):785–792.

53. Baylon EG, Crowder HA, Gold GE, Levenston ME. Non-ionic CT contrast solutions rapidly alter bovine cartilage and meniscus mechanics. Osteoarthritis and Cartilage. 2020 sep;28(9):1286–1297.

54. Chin HC, Moeini M, Quinn TM. Solute transport across the articular surface of injured cartilage. Archives of Biochemistry and Biophysics. 2013;535(2):241–247.

55. Bajpayee AG, Wong CR, Bawendi MG, Frank EH, Grodzinsky AJ. Avidin as a model for charge driven transport into cartilage and drug delivery for treating early stage post-traumatic osteoarthritis. Biomaterials. 2014;35(1):538–549.

56. Meganck JA, Kozloff KM, Thornton MM, Broski SM, Goldstein SA. Beam hardening artifacts in micro-computed tomography scanning can be reduced by X-ray beam filtration and the resulting images can be used to accurately measure BMD. Bone. 2009;45(6):1104–1116.

57. Edwards SH, Cake MA, Spoelstra G, Read RA. Biodistribution and clearance of intra-articular liposomes in a large animal model using a radiographic marker. Journal of Liposome Research. 2007;17(3-4):249–261.

58. Owen S, Francis H, Roberts M. Disappearance kinetics of solutes from synovial fluid after intra-articular injection. British Journal of Clinical Pharmacology. 1994;38(4):349–355.

59. Edwards SH. Intra-articular drug delivery: the challenge to extend drug residence time within the joint. The Veterinary Journal. 2011;190(1):15–21.

60. Leddy HA, Guilak F. Site-specific molecular diffusion in articular cartilage measured using fluorescence recovery after photobleaching. Annals of Biomedical Engineering. 2003;31(7):753–760.

61. Leddy HA, Haider MA, Guilak F. Diffusional anisotropy in collagenous tissues: fluorescence imaging of continuous point photobleaching. Biophysical Journal. 2006;91(1):311–316.

62. DiDomenico CD, Goodearl A, Yarilina A, Sun V, Mitra S, Sterman AS, et al. The effect of antibody size and mechanical loading on solute diffusion through the articular surface of cartilage. Journal of Biomechanical Engineering. 2017 sep;139(9). Available from: http://dx.doi.org/10.1115/1.4037202.

63. Pouran B, Arbabi V, Bajpayee AG, van Tiel J, Töyräs J, Jurvelin JS, et al. Multi-scale imaging techniques to investigate solute transport across articular cartilage. Journal of Biomechanics. 2018;78:10–20.

